# Boosting subdominant neutralizing antibody responses with a computationally designed epitope-focused immunogen

**DOI:** 10.1101/430157

**Authors:** F Sesterhenn, M Galloux, SS Vollers, L Csepregi, C Yang, D Descamps, J Bonet, S Friedensohn, P Gainza, P Corthésy, M Chen, S Rosset, MA Rameix-Welti, JF Eléouët, ST Reddy, BS Graham, S Riffault, BE Correia

## Abstract

Throughout the last decades, vaccination has been key to prevent and eradicate infectious diseases. However, many pathogens (e.g. respiratory syncytial virus (RSV), influenza, dengue and others) have resisted vaccine development efforts, largely due to the failure to induce potent antibody responses targeting conserved epitopes. Deep profiling of human B-cells often reveals potent neutralizing antibodies that emerge from natural infection, but these specificities are generally subdominant (i.e., are present in low titers). A major challenge for next-generation vaccines is to overcome established immunodominance hierarchies and focus antibody responses on crucial neutralization epitopes. Here, we show that a computationally designed epitope-focused immunogen presenting a single RSV neutralization epitope elicits superior epitope-specific responses compared to the viral fusion protein. In addition, the epitope-focused immunogen efficiently boosts antibodies targeting the Palivizumab epitope, resulting in enhanced neutralization. Overall, we show that epitope-focused immunogens can boost subdominant neutralizing antibody responses *in vivo* and reshape established antibody hierarchies.

## Introduction

The development of vaccines has proven to be one of the most successful medical interventions to reduce the burden of infectious diseases (*1*), and their correlate of protection is the induction of neutralizing antibodies (nAbs) that block infection (*2*).

In recent years, advances in high-throughput B-cell technologies have revealed a plethora of potent nAbs for different pathogens which have resisted the traditional means of vaccine development for several decades, including HIV-1 (*3*), influenza (*4*), respiratory syncytial virus (RSV) (*5, 6*), zika (*7, 8*), dengue (*9*) and others (*10-12*). A major target of these nAb responses is the pathogens fusion protein, which drives the viral and host cell membrane fusion while undergoing a conformational rearrangement from a prefusion to a postfusion state (*13*). Many of these nAbs have been structurally characterized in complex with their target, unveiling the atomic details of neutralization epitopes (*7, 14, 15*). Together, these studies have provided comprehensive antigenic maps of the viral fusion proteins which delineate epitopes susceptible to antibody-mediated neutralization and provide a roadmap for rational and structure-based vaccine design approaches.

The conceptual framework to leverage neutralizing antibody-defined epitopes for vaccine development is commonly referred to as reverse vaccinology (*16, 17*). Although reverse vaccinology-inspired approaches have yielded a number of exciting advances in the last decade, the design of immunogens that elicit such focused antibody responses remains challenging. Successful examples of structure-based immunogen design approaches include conformational stabilization of RSVF in its prefusion state, which induces superior serum neutralization titers when compared to immunization with F in the postfusion conformation (*18*). In the case of influenza, several epitopes targeted by broadly neutralizing antibodies (bnAbs) were identified within the hemagglutinin (HA) stem domain, and an HA stem-only immunogen elicited a broader neutralizing antibody response than full-length HA (*19, 20*). Commonly, these approaches have aimed to focus antibody responses on specific conformations or subdomains of viral proteins. In a more aggressive approach, Correia et al. (*21*) computationally designed a synthetic immunogen presenting the RSV antigenic site II (Figure 1a), and provided a proof-of-principle for the induction of site specific, RSV neutralizing antibodies using a synthetic immunogen.

**Figure 1:**
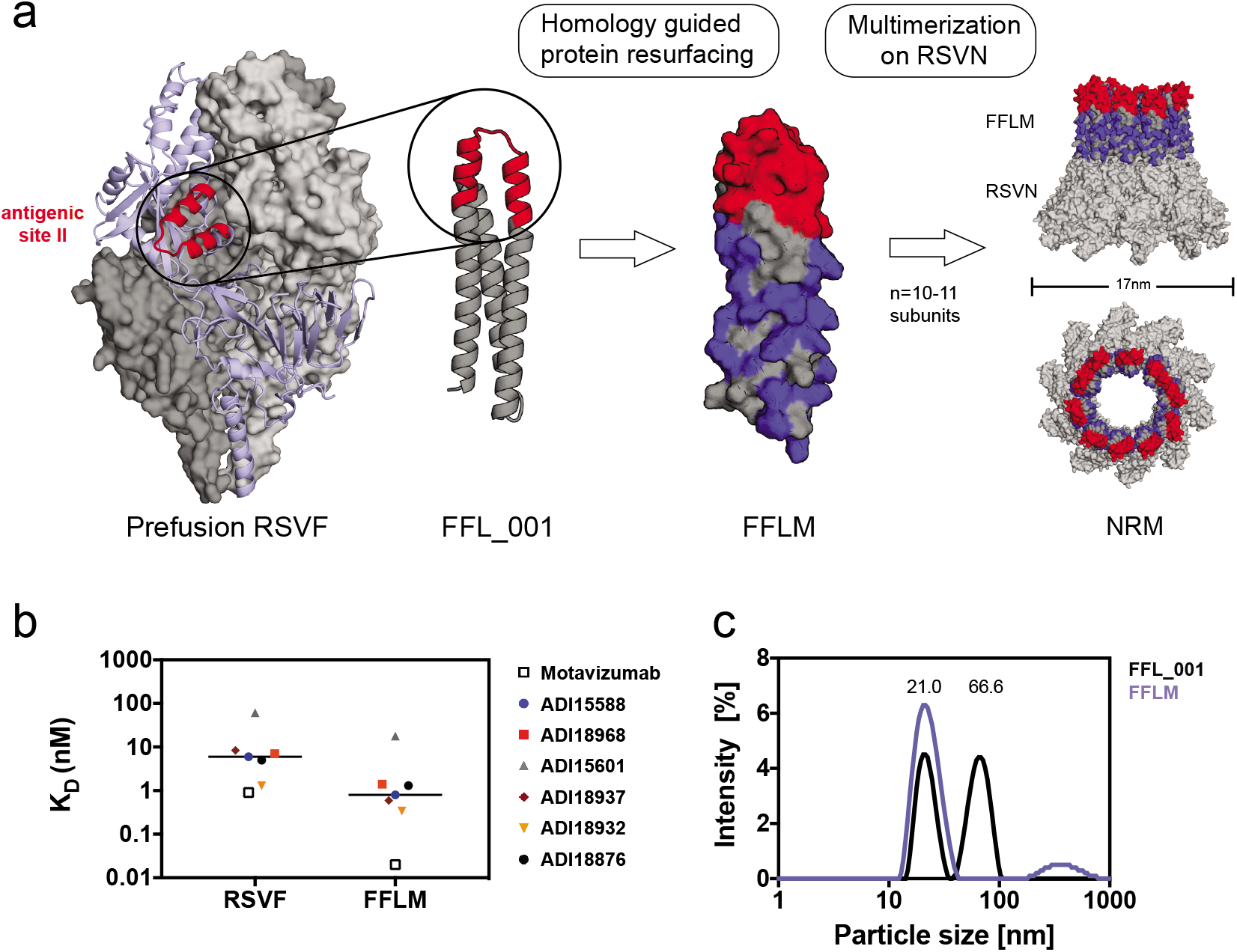
Design of an RSV-based nanoparticle displaying a site II epitope-focused immunogen. **a)**Structural model of the prefusion RSVF trimer (PDBID: 4JHW), with two subunits shown as a grey surface and one subunit shown as light blue cartoon representation with the epitope targeted by Palivizumab (antigenic site II) highlighted in red. FFL_001 was previously designed to present the site II epitope in a computationally designed scaffold. FFLM was designed by evolution-guided resurfacing, where changes in amino acid identity are highlighted in blue. FFLM was genetically fused to the N-terminus of the RSV nucleoprotein (RSVN), resulting in a high-density array of the epitope-scaffold, as shown by the structural model (based on PDBID: 2WJ8). **b)** Binding affinities of site II-specific human nAbs measured by SPR. KDs were measured with RSVF/FFLM immobilized as ligand and antibody fabs as analyte. nM = nanomolar. **c)** Dynamic light scattering (DLS) profiles for FFL_001 and FFLM fused to RSVN. The FFL_001-RSVN fusion protein formed higher-order oligomers in solution (66.6 nm of median diameter), whereas the resurfaced FFLM-RSVN fusion protein (NRM) was monodisperse with a median diameter of 21 nm. nm = nanometer.

The absence of a potent and long-lasting immune response upon natural infection is a major challenge associated with RSV, influenza virus and other pathogens. While single exposure to pathogens like poliovirus confers life-long immunity, RSV, influenza and other pathogens have developed mechanisms to subvert the development of a durable and potent neutralizing antibody response, thereby allowing such pathogens to infect humans repeatedly throughout their lives (*22*). One of the major factors hindering the induction of long-lasting protection after the first infection is related to the antibody specificities induced. Upon exposure to a pathogen, such as influenza, the human antibody responses predominantly target strain-specific antigenic sites, whereas potent bnAbs are subdominant (*23*). This phenomenon is generally referred to as B-cell immunodominance, which describes the unbalanced immunogenicity of certain antigenic sites within an antigen, favoring strain-specific, variable, non-neutralizing epitopes to the detriment of conserved, neutralization-sensitive epitopes (*24*). The factors that determine the antigenicity of specific epitopes remain unclear, making the categorization of immunodominant and subdominant epitopes an empirical classification based on serological analysis. Importantly, the presence of high levels of antibodies directed against immunodominant epitopes can sterically mask surrounding subdominant epitopes that may be targeted by bnAbs, preventing the immune system from mounting productive antibody responses against subdominant epitopes, and potentially limiting vaccination efficacy (*23-26*).

The immunodominance hierarchy is established within the germinal center, where B-cells undergo a binding affinity-based competition for available antigen and subsequently initiate a clonal expansion stage, ultimately becoming long-lived plasma cells or memory B-cells (*27*). Controlling this competition and driving antibody responses towards the increased recognition of subdominant, neutralizing epitopes is of primary importance to enable development of novel vaccines to fight pathogens which have resisted traditional strategies. One of the few strategies to guide antibody maturation was tested in the HIV field and is referred to as germline targeting, which relies upon the activation and expansion of rare but specific B-cell lineages in naïve individuals (*28, 29*). In contrast, under conditions of pre-existing immunity acquired during natural infection or previous vaccination, the challenge is to manipulate already established B-cell immunodominance hierarchies and reshape serum antibody responses towards desired specificities. In an indirect approach towards increasing subdominant B-cell populations, Silva et al. (*30*) have shown that the targeted suppression of immunodominant clones during an active germinal center reaction can allow subdominant B-cell populations to overtake the germinal center response. Other approaches have used heterologous prime-boost immunization regimens with either alternative viral strains or rationally modified versions of the priming immunogen (*31-34*) in order to steer antibody responses towards more conserved domains. However, leveraging structural information of defined neutralization epitopes to guide bulk antibody responses towards specific, well-characterized single epitopes remains an unmet challenge.

Here, we investigate whether, under conditions of pre-existing immunity, a computationally designed immunogen presenting a single epitope is able to reshape serum antibody responses towards increased recognition of a specific neutralizing epitope. To mimic a scenario of pre-existing immunity against a relevant pathogen, we immunized mice with a prefusion-stabilized version of RSVF, and found that antibody titers against RSV antigenic site II were present in very low levels, i.e. a subdominant epitope-specific response was elicited. Based on a previously developed epitope-focused immunogen for RSV site II (FFL_001) (*21*), we engineered an optimized nanoparticle presenting this immunogen, and investigated the potential of a rationally designed epitope-focused immunogen to boost these subdominant levels of site-specific antibodies.

We show that multivalent presentation of a designed epitope-focused immunogen elicits superior levels of epitope-specific antibodies compared to prefusion RSVF in naïve mice, indicating that the subdominance of a particular epitope can be altered through its presentation in a distinct molecular context. Repeated immunizations with RSVF failed to increase site II-specific antibodies, and instead further diluted site II specific responses. In contrast, heterologous boosts with an epitope-scaffold nanoparticle enhanced serum responses towards the subdominant site II epitope, and the boosted antibodies neutralized RSV *in vitro*. For the first time, we provide compelling evidence that synthetic immunogens comprising a single epitope can efficiently redirect specificities in bulk antibody responses *in vivo*. and enhance subdominant neutralizing antibody responses. Such strategy may present an important alternative for pathogens where future vaccines are required to reshape pre-existing immunity and elicit finely tuned antibody specificities.

## Results

### Design of an RSV-based nanoparticle displaying a site II epitope-focused immunogen

In a previous study, a computationally-designed, RSV site II epitope-scaffold nanoparticle was shown to elicit serum neutralization activity in non-human primates (NHPs) (*21*). Despite the fact that very potent monoclonal antibodies were isolated from the immunized NHPs, the neutralization potency at serum level was modest, indicating low titers of the potent antibodies. Therefore, our first aim was to take the best previously tested immunogen (FFL_001) and further optimize the immunogen delivery and immunization conditions to maximize the induction of site II-specific antibodies. A comparative study of four different adjuvants revealed that Alhydrogel^®^, an adjuvant approved for human use, yielded highest overall immunogenicity and elicited antibodies cross-reactive with prefusion RSVF in four out of five mice (Supplementary Fig.1).

Next, we sought to develop an improved, easily produced nanoparticle to multimerize the epitope-scaffold for efficient B-cell receptor crosslinking. Previously, Correia et al. (*21*) employed a chemical conjugation strategy of FFL_001 to a Hepatitis-B core antigen based nanoparticle, which resulted in a difficult construct with a laborious purification process. Recently, several studies have reported the use of RSV nucleoprotein (RSVN) as a nanoparticle platform for immunogen presentation (*35, 36*). When expressed in *E. coli*., RSVN forms nanorings, 17 nm in diameter, containing 10 or 11 RSVN protomers (*37*). We reasoned that RSVN would be an ideal particle platform to multimerize an RSV epitope-scaffold, as RSVN contains strong, RSV-directed T-cell epitopes (*36*). However, our initial attempts to genetically fuse FFL_001 to RSVN yielded poorly soluble proteins that rapidly aggregated after purification. We therefore employed structure-based protein resurfacing (*38*), attempting to improve the solubility of this site II epitope-scaffold when arrayed in high density on RSVN. To guide our resurfacing design process, we leveraged information from a sequence homolog of the ribosomal recycling factor (PDB: 1ISE), the structural template originally used to design FFL_001. Based on a sequence alignment of the mouse homolog (NCBI reference: NP_080698.1) and FFL_001, we exchanged the FFL_001 amino acids for the mouse sequence homolog and used Rosetta Fixed Backbone Design (*39*) to ensure that the mutations were not energetically unfavorable, resulting in 38 amino acid substitutions (34.2% overall). We named this variant FFLM, whose expression yields in *E. coli* showed a five-fold increase when compared to FFL_001, and it was confirmed to be monomeric in solution (Supplementary Fig.2).

To confirm that the resurfacing did not alter the epitope integrity, we measured the binding affinities of FFLM to Motavizumab, a high-affinity variant of Palivizumab (*40*), and to a panel of human site II nAbs previously isolated (*5*) using surface plasmon resonance (SPR). All antibodies bound with high affinity to FFLM, indicating broad reactivity of this immunogen with a diverse panel of human nAbs (Figure 1b). Interestingly, the tested nAbs showed approximately one order of magnitude higher affinity to the epitope-scaffold as compared to the latest version of prefusion RSVF, originally called DS2 (*41*), suggesting that the epitope is properly presented and likely further stabilized in a relevant conformation.

Importantly, the FFLM-RSVN fusion protein expressed with high yields in *E. coli*. (>10 mg/liter), forming a nanoring particle, dubbed NRM, that was monodisperse in solution with a diameter of approximately 21 nm (Figure 1c). Although we cannot fully rationalize the factors that contributed to the solubility improvement upon multimerization, our strategy to transplant surface residues from a sequence homolog to synthetic proteins may prove useful to enhance the solubility of other computationally designed proteins.

### NRM enhances the induction of site Il-specific antibodies

We next tested the immunogenicity of the site II scaffold nanoring and its ability to elicit site II-specific antibodies. Three groups of ten mice were subjected to three immunizations with 10 μg of NRM, monomeric FFLM and prefusion RSVF (*41*), which is currently the leading immunogen for an RSV vaccine (Figure 2a). Based on the results of our adjuvant screen (Supplementary Fig.1), all the immunogens were formulated in Alhydrogel, an adjuvant approved for human use. As compared to FFLM, NRM showed a higher overall immunogenicity (directed both against RSVN and FFLM) (Figure 2b).

**Figure 2:**
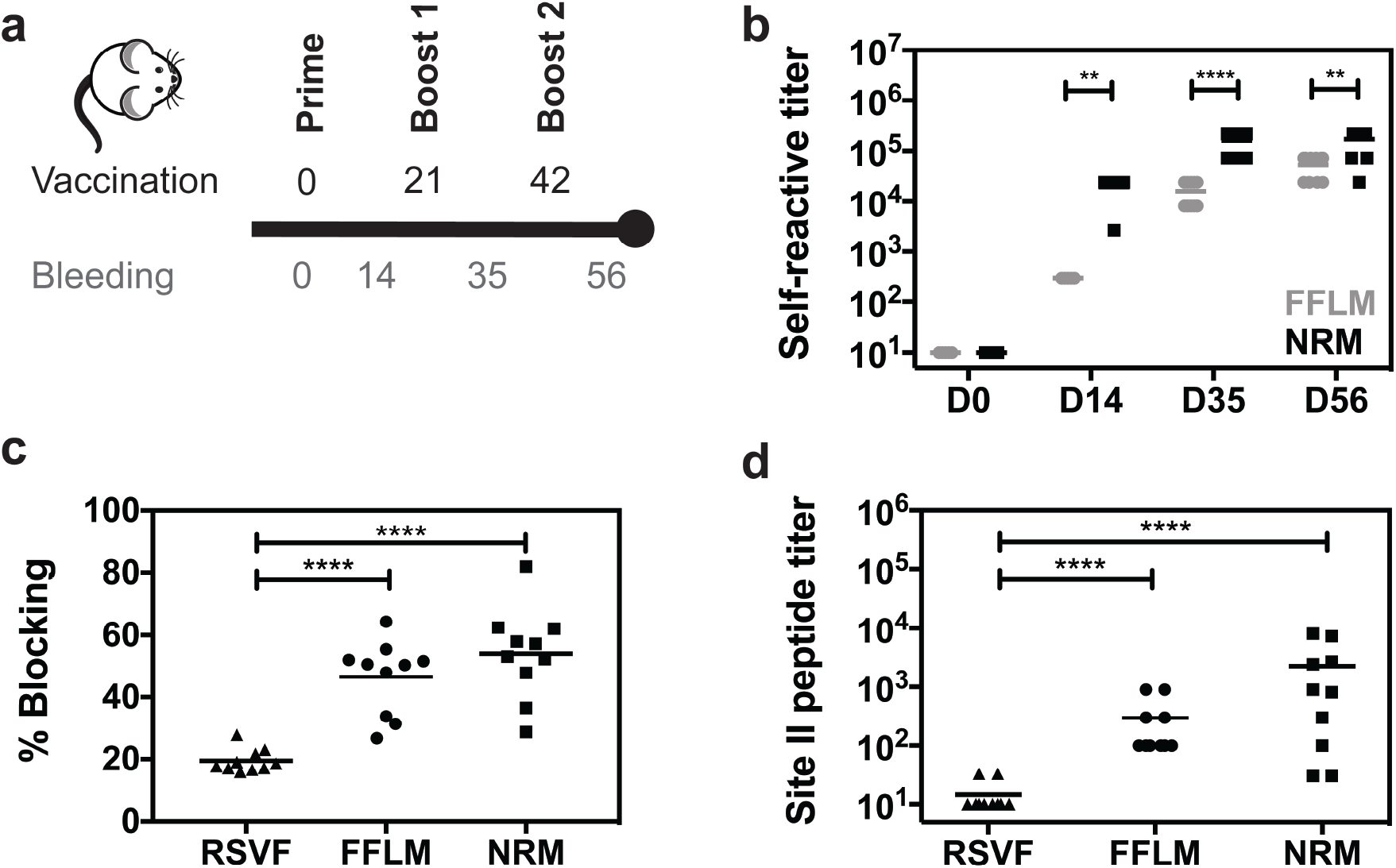
Immunogenicity and quantification of site II-specific antibody responses. **a**) Immunization scheme. Balb/c mice were immunized three times on days 0, 21 and 42, and blood was drawn 14 days after each vaccination. **b)** Serum antibody titers elicited by FFLM and NRM at different timepoints measured by ELISA against the respective immunogen. NRM shows significantly increased immunogenicity at day 14, 35 and 56 relative to FFLM. **c)** SPR competition assay with Motavizumab. Day 56 sera of mice immunized with RSVF, FFLM or NRM was diluted 1:100 and SPR response units (RU) were measured on sensor chip surfaces containing the respective immunogen. Motavizumab binding sites were then blocked by saturating amounts of Motavizumab, and the residual serum response was measured to calculate the serum fraction competed by Motavizumab binding. Mice immunized with FFLM or NRM show significantly higher levels of serum antibodies that are competed by Motavizumab binding. **d)** Site II-specific serum titers at day 56 from mice immunized with RSVF, FFLM and NRM, measured by ELISA against site II peptide. Three immunizations with prefusion RSVF elicited low levels of site II-specific antibodies, whereas FFLM and NRM vaccinations yielded significantly higher peptide-specific serum titers. Data shown are derived from at least two independent experiments, each sample assayed in duplicate. Statistical comparisons were calculated using two-tailed Mann-Whitney U tests. ** indicates p < 0.01, *** indicates p < 0.0001, **** p < 0.0001.

A key aspect of epitope-focused vaccines is to understand how much of the antibody response targets the viral epitope presented to the immune system. Therefore, we sought to measure the site II-specific antibody titers elicited by NRM and FFLM and compare these epitope-specific antibody responses to those elicited by prefusion RSVF. Using an SPR competition assay to measure site II-specific antibodies in sera (described in methods), we observed that NRM elicited site II-specific antibody responses superior to those elicited by RSVF (Figure 2c). This was surprising, given that the ratio of site II epitope surface area to overall immunogen surface is similar in both NRM and RSVF (Supplementary Fig.2). To confirm this finding through a direct binding assay rather than a competitive format, we measured the binding levels of sera to the site II epitope in a peptide ELISA, where the site II peptide was immobilized on a streptavidin-coated surface. Consistent with the previous experiment, we found that NRM elicited two orders of magnitude higher site II-specific responses than RSVF (Figure 2d). Together, we concluded that an epitope-focused immunogen, despite similar molecular surface area, can elicit substantially higher levels of site-specific antibodies compared to a viral fusion protein.

### NRM induces low levels of RSVF cross-reactive antibodies with low neutralization potency

Given the substantial site Il-specific serum titers elicited by NRM in mice, we investigated whether these antibodies cross-reacted with prefusion RSVF and were sufficient to neutralize RSV *in vitro*.

Following three immunizations with NRM, all the mice (n=10) developed detectable serum cross-reactivity with prefusion RSVF (mean serum titer = 980) (Figure 3a). Unsurprisingly, the overall quantity of prefusion RSVF cross-reactive antibodies elicited by immunization with an immunogen presenting a single epitope is more than two orders of magnitude lower than those of mice immunized with prefusion RSVF, which comprises at least six antigenic sites (*5*). Similarly, a B-cell ELISpot revealed that NRM-immunized mice presented prefusion RSVF-reactive antibody secreting cells, but their frequency was approximately one order of magnitude lower than upon immunization with prefusion RSVF (Figure 3c).

**Figure 3:**
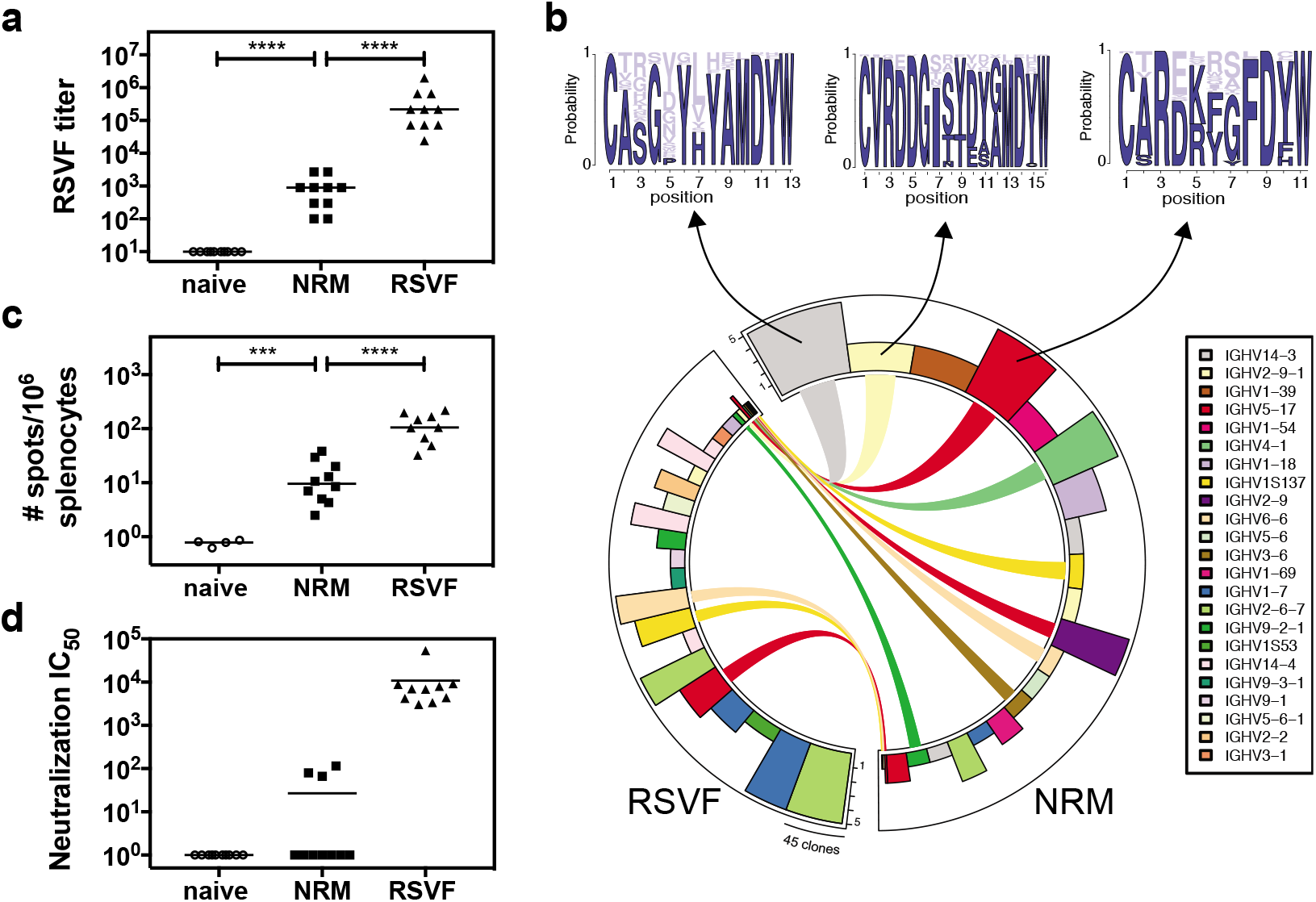
RSVF cross-reactivity and serum neutralization. **a)** NRM elicits prefusion RSVF cross-reactive antibodies, which are two orders of magnitude lower compared to prefusion RSVF immunization. Mice immunized only with adjuvant (naïve) do not show RSVF cross-reactivity.**b)** Next-generation sequencing of antibody repertoire. Antibody variable heavy chains of mice immunized with RSVF or NRM (5 mice per cohort) were sequenced and grouped into clonotypes. Circos plot showing the 20 most expanded clonotypes from both cohorts, with identical clonotypes connected. Height of bars indicates number of mice that showed the respective clonotype, width represents the clonal expansion within a clonotype (i.e. the number of clones grouped into the respective clonotype). Three clonotypes that occurred both in the RSVF and the NRM cohort, but were expanded within the NRM cohort were analyzed for their HCDR3 sequence profile, as shown by sequence logo plots (top). Dark blue color represents amino acid identities that occurred in RSVF cohort, light blue color represents amino acids uniquely found following NRM immunization. The frequency of each amino acid in the NRM cohort is indicated by the size of the letter. **c)** B-cell ELISpot of mouse splenocytes to quantify prefusion RSVF-specific antibody secreting cells (ASC). Number of ASCs per 10^6^ splenocytes that secrete prefusion RSVF-specific antibodies following three immunizations with adjuvant only (naïve), NRM or prefusion RSVF **d)** RSV neutralizing activity of mouse sera from day 56 shown as neutralization IC_50_. Three out of ten mice immunized with NRM showed detectable RSV neutralizing activity, whereas all mice immunized with prefusion RSVF neutralized RSV (mean IC_50_ = 10,827). Data shown are from one out of two independent experiments. Statistical comparisons were calculated using two-tailed Mann-Whitney U tests. *** indicates p < 0.001, **** indicates p < 0.0001.

The major determinant for antibody specificity is attributed to the heavy chain CDR3 region (HCDR3) (*42*). While for certain classes of nAbs, the antibody lineages and their sequence features are well-defined (e.g. HIV neutralizing VRC01 class antibodies (*43*), or RSV neutralizing MPE8-like antibodies (*44*)), antibodies targeting RSV antigenic site II seem to be derived from diverse precursors and do not show HCDR3 sequence convergence in humans (*5*). While we did not expect to find dominant lineages or HCDR3 sequence patterns in mice, we used next-generation antibody repertoire sequencing (*45*) to ask whether NRM could elicit antibodies with similar sequence signatures to those elicited by prefusion RSVF. Indeed, we found 300 clonotypes, defined as antibodies derived from the same VH gene with the same HCDR3 length and 80% sequence similarity, that overlapped between NRM and the prefusion RSVF immunized cohort, suggesting that at the molecular level, relevant antibody lineages can be activated with the NRM immunogen (Supplementary Figure 3). Notably, nine out of the 20 most expanded clonotypes in the NRM cohort were also present in mice immunized with prefusion RSVF, albeit not as expanded (Figure 3b). This finding might reflect the enrichment of site II specific antibodies in the NRM cohort (Figure 2d).

We further investigated whether these low levels of prefusion RSVF-binding antibodies were sufficient to neutralize RSV *in vitro* While three immunizations with prefusion RSVF elicited potent RSV-neutralizing serum titers (mean IC_50_= 10,827), for NRM we only detected low levels of RSV-neutralizing serum activity in three out of ten mice (Figure 3d). This result is consistent with Correia *et al*. (*21*), who observed no serum neutralization in mice, but succeeded in inducing nAbs in NHPs with prior RSV seronegativity.

Altogether, we concluded that despite NRM’s superior potential to induce high levels of site II-specific antibodies, the majority of antibodies activated from the naïve repertoire is not functional for RSV neutralization. A potential explanation, stemming from structural comparison between the epitope-focused immunogen (FFLM) and RSVF, is that although epitope-specific antibodies are abundantly elicited by NRM, these antibodies do not recognize the site II epitope in its native RSVF quaternary environment in the prefusion conformation, or on virions in sufficient amounts and with high enough affinity to potently neutralize RSV.

### NRM boosts site Il-specific antibodies under conditions of pre-existing immunity

While vaccination studies in naïve animal models are an important first step to validate novel immunogens, previous studies (*21*) and results presented here imply that epitope-scaffolds may not be able to elicit robust RSV neutralizing serum activity from a naïve antibody repertoire. However, given the high affinity of the epitope-scaffold towards a panel of site II-specific nAbs, together with the ability to elicit high titers of site II-specific antibodies *in vivo*, we hypothesized that such an epitope focused immunogen could be efficient in recalling site II-specific B-cells in a scenario of pre-existing immunity, thereby achieving an enhanced site-specific neutralization response.

Our initial immunization studies with prefusion RSVF showed that site II-specific responses were subdominant (Figure 2c and 2d). Given that subdominance is a common immunological phenotype for many of the neutralization epitopes that are relevant for vaccine development (*46*), we sought to test if NRM could boost subdominant antibody lineages that should ultimately be functional and recognize the epitope in the tertiary environment of the viral protein. To test this hypothesis, we designed a mouse immunization experiment with three cohorts, as outlined in Figure 4a. Following a priming immunization with RSVF, cohort (1) was boosted with adjuvant only (“prime only”), cohort (2) received two boosting immunizations with prefusion RSVF (“homologous boost”), and cohort (3) received two boosts with NRM (“heterologous boost”).

**Figure 4:**
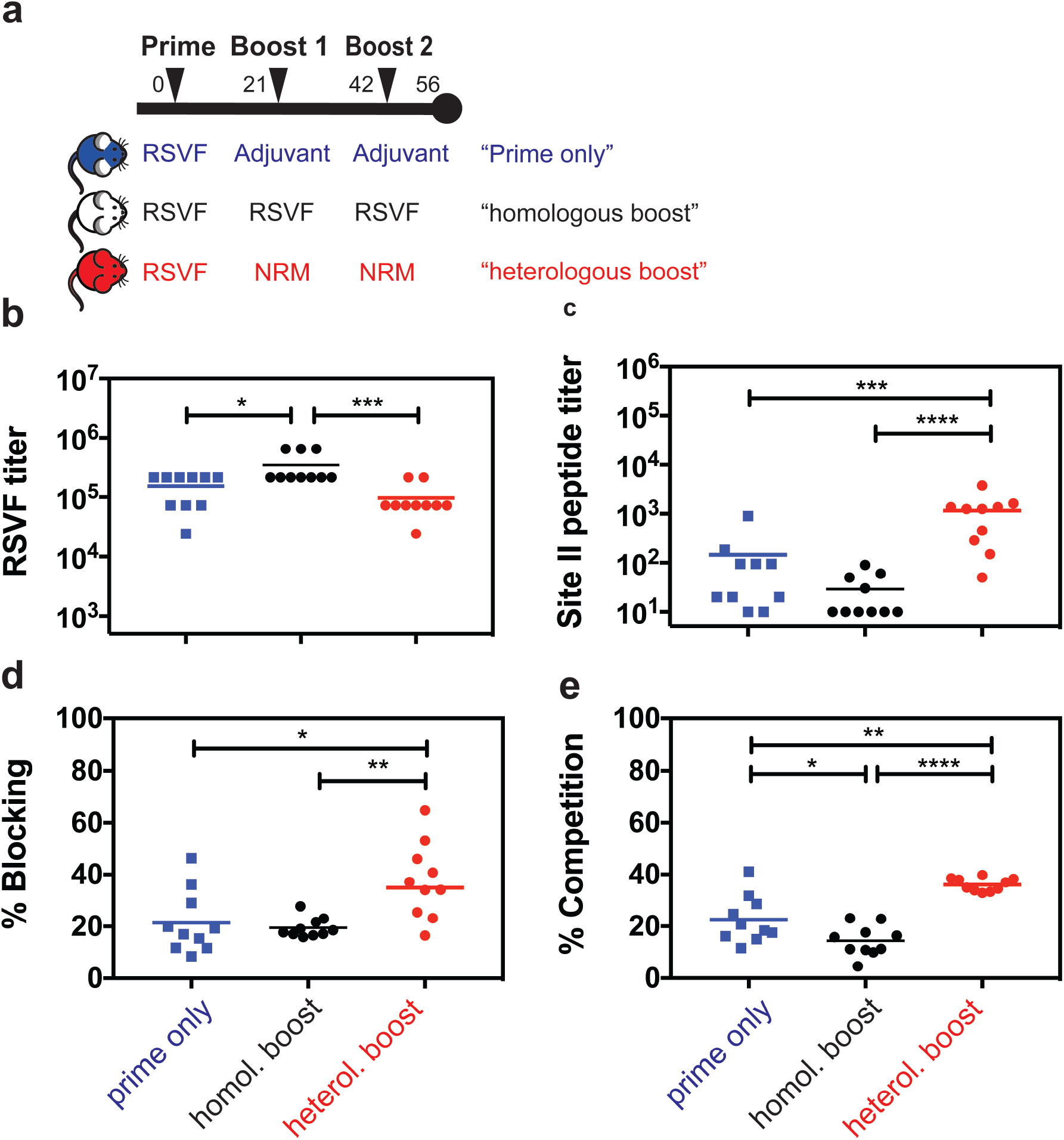
Heterologous prime boost reshapes antibody responses enhancing levels of site II specific antibodies. **a)** Heterologous prime-boost study groups. Three mouse cohorts were immunized with either 1x RSVF (“prime only”), 3x RSVF (“homologous boost”) or 1x RSVF followed by two boosts with NRM (“heterologous boost”). **b)** Antibody titers directed against prefusion RSVF. Mice receiving homologous boosting immunizations show slightly higher RSVF-specific serum titers compared to the prime only cohort, whereas heterologous boosting yielded statistically comparable titers to the prime only group. The difference between the homologous and heterologous boost cohorts was statistically significant. **c)** Site II-specific titers measured by ELISA showed that the heterologous boost significantly increases site II-specific titers compared to both prime and homologous boost groups. Albeit not statistically significant (p = 0.06), mice receiving a homologous boost had lower levels of site II-specific antibodies compared to prime only group. **d)** SPR competition assay with Motavizumab on a prefusion RSVF-coated sensor chip. Sera from indicated groups were diluted 1:100 and RSVF binding responses were quantified. Site II was then blocked with Motavizumab, and the remaining serum response quantified. The heterologous boost induced a significantly higher fraction of site II-directed antibodies competed with Motavizumab for RSVF binding, as compared to both prime only and homologous boost groups). **e)** Quantification of site II-specific responses in a competition ELISA. Binding was measured against prefusion RSVF, and the Area Under the Curve (AUC) was calculated in presence of NRM competitor, normalized to the AUC in the presence of RSVN as a control competitor. Compared to the prime only group, the homologous boost resulted in significantly lower site II-specific serum titers, confirming the trend observed in c). The heterologous boost increased the fraction of site II-targeting antibodies within the pool of prefusion RSVF-specific antibodies compared to both control groups. Data presented are from at least two independent experiments, with each sample assayed in duplicates. Statistical comparisons were calculated using two-tailed Mann-Whitney U tests. * indicates p<0.05, ** indicates p < 0.01, *** indicates p < 0.0001, **** p < 0.0001.

A comparison between prefusion RSVF immunized groups prime only and homologous boost revealed that the two additional boosting immunizations with RSVF only slightly increased overall titers of prefusion RSVF-specific antibodies (p = 0.02), indicating that a single immunization with adjuvanted RSVF is sufficient to induce close to maximal serum titers against RSVF (Figure 4b). Following the heterologous boost with NRM, overall RSVF specific antibody titers remained statistically comparable to the prime only group (p = 0.22).

Next, we quantified the site II-specific endpoint serum titers in a peptide ELISA format (Figure 4c). Interestingly, the homologous boost with prefusion RSVF failed to increase site II-specific antibody levels, reducing the responses directed to site II to the lower limit of detection by ELISA. This result is yet another example of the underlying complexity inherent to the fine specificity of antibody responses elicited by immunogens and how important specificities can be dampened throughout the development of an antibody response. In contrast to the homologous boost, the heterologous boost with NRM significantly increased site II peptide-specific serum titers (p < 0.0001).

In order to understand whether this increase relied at least partially on an actual recall of antibodies primed by RSVF, or rather on an independent antibody response irrelevant for RSVF binding and RSV neutralization, we dissected the epitope specificity within the RSVF-specific serum response. In an SPR competition assay, a significantly higher fraction (p = 0.02) of prefusion RSVF-reactive antibodies were competed by Motavizumab in mouse sera primed with prefusion RSVF and boosted with NRM (mean % competition = 37.5 ± 14.5%), as compared to mice immunized once or three times with prefusion RSVF (21.5% ± 12.1% or 19.5% ± 3.7%, respectively) (Figure 4d). Similarly, a competition ELISA revealed that a significantly larger fraction of overall RSVF reactivity was attributed to site II-specific antibodies upon heterologous boost, as compared to both control groups (36.1% ± 2.5% versus 22.6% ± 9.1% or 14.4% ±5.9%, respectively, p = 0.002 and p < 0.0001). In contrast, site II-specific antibodies were significantly higher in mice that received only one as opposed to three RSVF immunizations, indicating that RSVF boosting immunizations further diluted site II-specific antibody titers (p = 0.03) (Figure 4e).

Together, we have shown that the serum antibody specificity can be steered towards a well-defined antigenic site by boosting pre-existing, subdominant antibody levels with an epitope focused immunogen. This is an important and distinctive feature of the epitope focused immunogen compared to an immunogen based on a viral protein (prefusion RSVF), which was shown to decrease already subdominant antibody responses under the same conditions. These results may have broad implications on strategies to control antibody fine specificities in vaccination schemes, both for RSV and other pathogens.

### Boosted antibodies neutralize RSV *in vitro*

The enhanced reactivity to site II observed in the heterologous prime-boost scheme led us to investigate if antibodies boosted by a synthetic immunogen were functionally relevant for virus neutralization. In bulk sera, we observed 2.3-fold higher serum neutralization titers in mice receiving a heterologous boost (mean IC_50_=7,654) compared to the prime only control group (mean IC_50_=3,275) (Figure 5a). While this increase in serum neutralization was not statistically significant, we next assessed if this increase in neutralization was driven by increased levels of epitope-specific antibodies. We observed that site II-directed antibody levels correlated with overall serum neutralization titers in the heterologous prime boost group (r^2^=0.76, p=0.0009) (Figure 5b), whereas prime only (r^2^=0.32, p=0.09) or animals receiving a homologous boost showed no such correlation (r^2^ = 0.18, p=0.22) (Supplementary Figure 4). To characterize the neutralizing serum activity dependence on antigenic site II, we pooled mouse sera within each cohort, enriched site II-specific antibodies and measured viral neutralization (see methods). Briefly, we incubated pooled sera from each group with streptavidin beads conjugated to biotin-labeled antigenic site II peptide, and eluted bound antibodies. To control for the quality of the enrichment protocol, we verified by ELISA that the column flow-through was depleted of site II-specific antibodies (Supplementary Figure 5).

**Figure 5.**
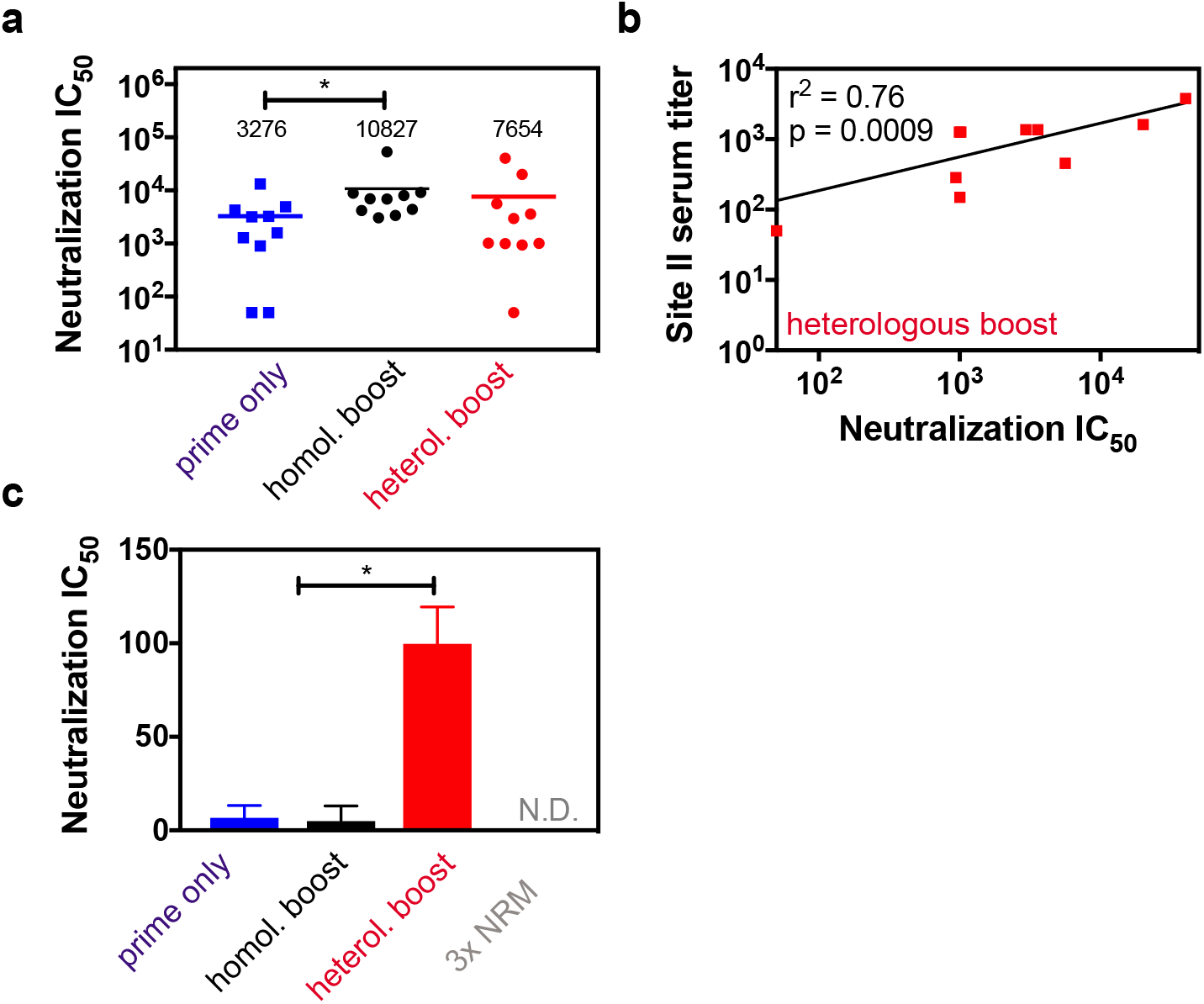
Boosted site II-specific antibodies are functional and mediate increased neutralization activity. **a)** *in vitro* RSV neutralization IC_50_ for each group. Compared to the prime only group, mice receiving a homologous boost showed increased RSV neutralization titers. On average, the heterologous boost yielded a 2.3-fold increase in serum neutralization titers compared to prime only, but these differences were statistically not significant when compared to either group. **b)** Correlation of site II-specific serum titer (measured by peptide ELISA) with RSV neutralization IC_50_ as determined for each mouse within the heterologous prime boost cohort. Correlations for control groups are shown in Supplementary Figure 4. Data represent the mean of two independent experiments, each measured in duplicate. Pearson correlation coefficient (r^2^) and p-value were calculated in GraphPad Prism. **c)** Site II-specific antibody fractionation revealed increased levels of nAbs. Site II-specific antibodies from mouse sera were enriched in an affinity purification. Those isolated from the heterologous boost group showed a 15-fold increase in RSV neutralizing activity compared to both control groups. No site II-mediated neutralization was detected for mice receiving three immunizations of NRM (N.D. = non-detectable). Data are presented from two independent experiments, and each sample was assayed in duplicate with additional controls shown in Supplementary Figure 5. Statistical comparisons were calculated using two-tailed Mann-Whitney U tests. * indicates p < 0.05.

Strikingly, mice receiving a heterologous boost showed a 15-fold increase in site II-mediated neutralization as compared to mice immunized once or three times with prefusion RSVF (Figure 5c). We performed the same experiment for mice immunized three times with NRM and did not detect any RSV neutralization in this format. This observation is consistent with the very low levels of bulk sera neutralization measured in this group, indicating that NRM can only boost nAbs under conditions of pre-existing immunity. In summary, we conclude that the heterologous boosting scheme with a single epitope immunogen enhanced subdominant neutralizing antibody responses directed against the antigenic site presented, and effectively redirected an antibody response *in vivo*.

## Discussion

Despite a rapid increase in our atomic-level understanding of antibody-antigen interactions for various pathogens, the translation of structural information into efficacious immunogens that elicit antibody responses specific to *bona fide* epitopes remains a key challenge for next-generation vaccine development.

Multiple strategies have been investigated to focus nAb responses on defined neutralization epitopes (*47*). Among them, epitope-scaffolds have been shown to elicit RSV site Il-specific, neutralizing antibody responses in naïve non-human primates. While the overall serum neutralization was modest, a monoclonal antibody induced by vaccination, showed superior neutralization potency to that of Palivizumab (*21*). However, a major limitation of epitope-scaffold immunogens (*48-50*) is that the quaternary environment of the epitope presented in the native viral protein is lost. Thus, the binding mode of a significant fraction of the elicited antibodies is likely incompatible with the epitope in its native environment. This observation is reinforced by our finding that although NRM elicited high serum levels of site II-directed antibodies, only residual neutralizing activity was observed in mice, which is consistent with previous studies using epitope-scaffolds (*50-52*). Together, these results highlight the limitations of synthetic scaffolds in an epitope-focused vaccine approach in naïve individuals.

However, our finding that a subdominant epitope (site II) in its native environment (prefusion RSVF) is readily targeted by the immune system when presented in a distinct molecular context (NRM), supported the potential use of synthetic immunogens to reshape antibody responses towards *bona fide* vaccine epitopes. Pre-existing immunity against a viral protein (RSVF, influenza HA or others), in which certain antibody specificities are subdominant, is a common scenario in humans that have encountered repeated natural infections throughout their life (*26, 53-55*). Therefore, a major challenge for vaccine development is to boost pre-existing, subdominant antibodies to enhance site-specific neutralization.

To date, boosting nAbs targeting specific epitopes under conditions of pre-existing immunity has been challenging. For instance, strong antibody responses against immunodominant epitopes can sterically mask the neutralization epitope, preventing the induction of a potent antibody response targeting the subdominant site (*23, 25, 26, 56*). Overcoming these established immunodominance hierarchies is complex, as such hierarchies seem to be impacted by multiple factors including serological antibody levels, their specificity, memory B-cell counts, adjuvants, and the immunization or infection route (*24*).

Heterologous prime-boost schemes are a promising strategy to guide the fine specificity of antibody responses and to focus these responses on vulnerable antigenic sites. Several vaccine studies have been conducted for influenza (*33, 34*), RSV (*31*) and HIV (*28*), where the heterologous immunogens were alternative strains or modified viral fusion proteins, but yet not as heterologous as a computationally designed epitope-scaffold. An important point to consider regarding immunogens based on modified viral proteins is whether immunodominant signatures remain, steering the antibody responses away from the target epitopes. While this scenario may not be fully absent in synthetic epitope-scaffolds, it is at least mitigated by the fact that the protein has not evolved under the pressure of escaping the immune system.

Our study demonstrates that a heterologous boosting immunogen with a single neutralization epitope, when optimally presented can enhance pre-existing, subdominant antibody responses targeting this epitope. The ability to narrowly focus antibody responses to a single epitope that mediates clinical protection, underlines the potential of rationally designed immunogens for vaccine development against elusive pathogens. In particular, our results demonstrate that albeit single-epitope immunogens may not be the most powerful to select functional antibodies from a naïve repertoire, they have a unique ability to boost neutralizing epitope-specific antibodies primed by a viral protein. Further studies in more relevant animal models will reveal if neutralizing antibodies primed by natural infection with RSV can also be boosted mimicking a more realistic vaccination scenario.

Given that the approach presented here is generalizable and that epitope-scaffold nanoparticles can be proven successful in boosting nAbs specific for other sites, this strategy holds great potential to tune levels of antibody specificities through heterologous prime boost vaccination schemes which are now frequently used in for challenging pathogens (*28, 33, 57*).

The original antigenic sin theory in the influenza field describes that the first viral exposure permanently shapes the antibody response, which causes individuals to respond to seasonal vaccines dependent on their immune history (*23, 58*). Seasonal vaccines generally fail to boost antibodies targeting broadly neutralization epitopes on the hemagglutinin stem region (*23*). Focusing antibody responses on these defined epitopes may remove the need for annual vaccine reformulation, and may also protect against emerging pandemic strains (*14, 46, 59, 60*). The influenza vaccine challenge seems particularly well suited to our approach considering that the human population has pre-existing immunity to influenza, including some subdominant bnAbs that seasonal vaccines fail to stimulate (*23*).

Lastly, vaccine development against antigenically related viruses such as zika and dengue could benefit of the approach presented here, as antibodies mounted against the envelope protein of a dengue subtype can facilitate infection with zika (*61*) or other dengue subtypes (*62*). A site conserved between all four dengue subtypes and zika envelope protein has been structurally characterized and suggested for the development of an epitope-focused immunogen (*7*).

When seeking to apply an immunofocusing strategy to other antigenic sites and pathogens, one challenge is the development of epitope-scaffolds stably presenting the epitope in a synthetic immunogen that is compatible with antibody binding. While the RSV antigenic site II is a structurally simple helix-turn-helix motif, many other identified neutralization epitopes comprise multiple, discontinuous segments. However, continuous advances in rational protein design techniques (*63*) will allow the design of more complex protein scaffolds to stabilize increasingly complex epitopes.

Altogether, we have shown how an optimized presentation of a computationally designed immunogen in an RSVN-based nanoparticle can reshape bulk serum responses and boost subdominant, neutralizing antibody responses *in vivo*. This is a distinctive feature compared to using prefusion RSVF as a boosting immunogen, and underscores how subdominant epitopes can be converted to immunodominant epitopes when presented in a different environment. We foresee the great promise of this strategy to overcome the challenge of boosting and focusing pre-existing immunity towards defined neutralization epitopes, potentially applicable to multiple pathogens.

## Methods

### Resurfacing

The previously published RSV site II epitope-scaffold (“FFL_001”) (*21*) was designed based on a crystal structure of a mutant of ribosome recycling factor from *E. coli* (PDB entry 1ISE. Using BLAST, we identified sequence homologs of 1ISE from eukaryotic organisms and created a multiple sequence alignment with clustal omega (CLUSTALO (1.2.1)) (*64*) of the mouse homolog sequence (NCBI reference NP_080698.1), 1ISE and FFL_001. Surface-exposed residues of FFL_001 were then mutated to the respective residue of the mouse homolog using the Rosetta fixed backbone design application (*39*), resulting in 38 surface mutations. Amino acid changes were verified to not impact overall Rosetta energy score term.

### Protein expression and purification

#### FFLM

DNA sequences of the epitope-scaffold designs were purchased from Genscript and cloned in pET29b, in frame with a C-terminal 6x His tag. The plasmid was transformed in *E. coli* BL21 (DE3) and grown in Terrific Broth supplemented with Kanamycin (50 μg/ml). Cultures were inoculated to an OD_600_ of 0.1 from an overnight culture and incubated at 37°C. After reaching OD_600_ of 0.6, expression was induced by the addition of 1 mM isopropyl-β-D-thiogalactoside (IPTG) and cells were incubated for further 4-5h at 37°C. Cell pellets were resuspended in lysis buffer (50 mM TRIS, pH 7.5, 500 mM NaCl, 5% Glycerol, 1 mg/ml lysozyme, 1 mM PMSF, 1 μg/ml DNase) and sonicated on ice for a total of 12 minutes, in intervals of 15 seconds sonication followed by a 45 seconds pause. Lysates were clarified by centrifugation (18,000 rpm, 20 minutes), sterile-filtered and purified using a His-Trap FF column on an Äkta pure system (GE healthcare). Bound proteins were eluted in buffer containing 50 mM Tris, 500 mM NaCl and 300 mM imidazole, pH 7.5. Concentrated proteins were further purified by size exclusion chromatography on a Superdex™ 75 300/10 (GE Healthcare) in PBS. Protein concentrations were determined via measuring the absorbance at 280 nm on a Nanodrop (Thermo Scientific). Proteins were concentrated by centrifugation (Millipore, #UFC900324) to 1 mg/ml, snap frozen in liquid nitrogen and stored at −80°C.

#### NRM

The full-length N gene (sequence derived from the human RSV strain Long, ATCC VR-26; GenBank accession number AY911262.1) was PCR amplified using the Phusion DNA polymerase (Thermo Scientific) and cloned into pET28a+ at NcoI-XhoI sites to obtain the pET-N plasmid. The sequence of FFLM was then PCR amplified and cloned into pET-N at NcoI site to the pET-NRM plasmid *E. coli* BL21 (DE3) bacteria were co-transformed with pGEX-PCT (*65*) and pET-FFLM-N plasmids and grown in LB medium containing ampicillin (100 μg/ml) and kanamycin (50 μg/ml). The same volume of LB medium was then added, and protein expression was induced by the addition of 0.33 mM IPTG to the medium. Bacteria were incubated for 15 h at 28°C and then harvested by centrifugation. For protein purification, bacterial pellets were resuspended in lysis buffer (50 mM Tris-HCl pH 7.8, 60 mM NaCl, 1 mM EDTA, 2 mM dithiothreitol, 0.2% Triton X-100, 1 mg/ml lysozyme) supplemented with a complete protease inhibitor cocktail (Roche), incubated for one hour on ice, and disrupted by sonication. The soluble fraction was collected by centrifugation at 4 °C for 30 min at 10,000 × g. Glutathione-Sepharose 4B beads (GE Healthcare) were added to clarify supernatants and incubated at 4°C for 15h. The beads were then washed one time in lysis buffer and two times in 20 mM Tris pH 8.5, 150 mM NaCl. To isolate NRM, beads containing bound complex were incubated with thrombine for 16 h at 20 °C. After cleavage of the GST tag, the supernantant was loaded onto a Sephacryl S-200 HR 16/30 column (GE Healthcare) and eluted in 20 mM Tris-HCl, 150 mM NaCl, pH 8.5.

#### Antibody variable fragments (Fabs)

For Fab expression, heavy and light chain DNA sequences were purchased from Twist Biosciences and cloned separately into the pHLSec mammalian expression vector (Addgene, #99845) using AgeI and XhoI restriction sites. Expression plasmids were pre-mixed in a 1:1 stoichiometric ratio, co-transfected into HEK293-F cells and cultured in FreeStyle™ medium (Gibco, #12338018). Supernatants were harvested after one week by centrifugation and purified using a kappa-select column (GE Healthcare). Elution of bound proteins was conducted using 0.1 M glycine buffer (pH 2.7) and eluates were immediately neutralized by the addition of 1 M Tris ethylamine (pH 9), followed by buffer exchange to PBS pH 7.4.

#### Respiratory Syncytial Virus Fusion protein (prefusion RSVF)

Protein sequence of prefusion RSVF corresponds to the sc9-10 DS-Cav1 A149C Y458C S46G E92D S215P K465Q variant designed by Joyce et al. (*41*), which we refer to as RSVF DS2.RSVF DS2 was codon optimized for mammalian expression and cloned into the pHCMV-1 vector together with two C-terminal Strep-Tag II and one 8x His tag. Plasmids were transfected in HEK293-F cells and cultured in FreeStyle™ medium. Supernatants were harvested one week after transfection and purified via Ni-NTA affinity chromatography. Bound protein was eluted using buffer containing 10 mM Tris, 500 mM NaCl and 300 mM Imidazole (pH 7.5), and eluate was further purified on a StrepTrap HP affinity column (GE Healthcare). Bound protein was eluted in 10mM Tris, 150 mM NaCl and 20 mM Desthiobiotin (Sigma), pH 8, and size excluded in PBS, pH 7.4, on a Superdex 200 Increase 10/300 GL column (GE Healthcare) to obtain trimeric RSVF.

### Affinity determination using Surface Plasmon Resonance

Surface Plasmon Resonance experiments were performed on a Biacore 8K at room temperature with HBS-EP+ running buffer (10 mM HEPES pH 7.4, 150 mM NaCl, 3 mM EDTA, 0.005 % v/v Surfactant P20) (GE Healthcare). Approximately 100 response units (RU) of FFLM were immobilized via amine coupling on a CM5 sensor chip (GE Healthcare). Serial dilutions of site II-specific antibody variable fragments (fabs) were injected as analyte at a flow rate of 30 μl/min with 120 seconds contact time. Following each injection cycle, ligand regeneration was performed using 0.1 M glycine, pH 2. Data analysis was performed using 1:1 Langmuir binding kinetic fits within the Biacore evaluation software (GE Healthcare).

### Mouse immunizations

All animal experiments were approved by the Veterinary Authority of the Canton of Vaud (Switzerland) according to Swiss regulations of animal welfare (animal protocol number 3074). Six-week-old, female Balb/c mice were ordered from Janvier labs and acclimatized for one week. Immunogens were thawed on ice and diluted in PBS pH 7.4 to a concentration of 0.2 mg/ml. The immunogens were then mixed with an equal volume of 2% Alhydrogel^®^ (Invivogen), resulting in a final Alhydrogel concentration of 1%. Other adjuvants were formulated according to manufacturer’s instructions. After mixing immunogens and adjuvants for one hour at 4°C, each mouse was injected with 100 μl, corresponding to 10 μg immunogen adsorbed to Alhydrogel. All immunizations were done subcutaneously, with no visible irritation around the injection site. Immunizations were performed on day 0, 21 and 42. 100-200 μl blood were drawn on day 0, 14, 35, and the maximum amount of blood (200-1000μl) was taken by cardiac puncture at day 56, when mice were sacrificed.

### Antigen ELISA

Nunc Medisorp plates (Thermo Scientific, # 467320) were coated overnight at 4°C with 100 μl of antigen (recombinant RSVF, FFLM and NRM) diluted in coating buffer (100 mM sodium bicarbonate, pH 9) at a final concentration of 0.5 μg/ml. For blocking, plates were incubated for two hours at room temperature with blocking buffer (PBS + 0.05% Tween 20 (PBST) supplemented with 5% skim milk powder (Sigma, #70166)). Mouse sera were serially diluted in blocking buffer and incubated for one hour at room temperature. Plates were washed five times with PBST before adding 100 μl of anti-mouse HRP-conjugated secondary antibody diluted at 1:1500 in blocking buffer (abcam, #ab99617). An additional five washes were performed before adding Pierce TMB substrate (Thermo Scientific, #34021). The reaction was stopped by adding 100 μl of 2M sulfuric acid, and absorbance at 450 nm was measured on a Tecan Safire 2 plate reader.

Each plate contained a standard curve of Motavizumab to normalize signals between different plates and experiments. Normalization was done in GraphPad Prism. The mean value was plotted for each cohort and statistical analysis was performed using GraphPad Prism.

### Competition ELISA

Prior to incubation with a coated antigen plate, sera were serially diluted in the presence of 100 μg/ml competitor antigen and incubated overnight at 4°C. ELISA curves of a positive control, Motavizumab, are shown in Supplementary Figure 6. Curves were plotted using GraphPad Prism, and the area under the curve (AUC) was calculated for the specific (NRM) and control (RSVN) competitor. % competition was calculated using the following formula (*66*):

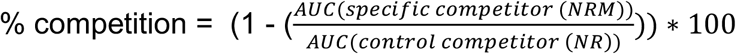

### Peptide sandwich ELISA

The antigenic site II was synthesized as peptide by JPT Peptide Technologies, Germany. The following sequence was synthesized and biotinylated at the N-terminus:

MLTNSELLSKINDMPITNDQKKLMSNNVQI

For ELISA analysis of peptide-reactive serum antibodies, Nunc MediSorp plates were coated with 5 μg/ml streptavidin (Thermo Scientific, #21122) for one hour at 37°C. Subsequently, ELISA plates were blocked as indicated above, followed by the addition of 2.4 μg/ml of the biotinylated site II peptide. Coupling was performed for one hour at room temperature. The subsequent steps were performed as described for the antigen ELISA.

### Serum competition using Surface Plasmon Resonance

Approximately 300 RU of antigen were immobilized via amine coupling on a CM5 chip. Mouse sera were diluted 1:100 in HBS-EP+ running buffer and flowed as analyte with a contact time of 120 seconds to obtain an initial response unit (RU_non-blocked surface_). The surface was regenerated using 50 mM NaOH. Sequentially, Motavizumab was injected four times at a concentration of 2 μM, leading to complete blocking of Motavizumab binding sites as confirmed by signal saturation. The same serum dilution was reinjected to determine the remaining response (RU_blocked surface_). The delta serum response (Δ *SR*) corresponds to the baseline-subtracted, maximum signal of the injected sera.

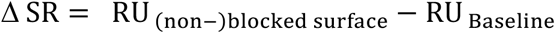

Percent blocking was calculated as follows:

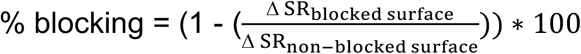

A schematic representation of the SPR experiment is shown in Supplementary Figure 7, and calculated blocking values are shown in Supplementary Table 1.

### Enzyme-linked immunospot assay (ELISPOT)

B-cell ELISPOT assays were performed using the Mouse IgG ELISpot HRP kit (Mabtech, #3825-2H) according to the manufacturer’s instructions. Briefly, mouse spleens were isolated, and pressed through a cell strainer (Corning, #352350) to obtain a single cell suspension. Splenocytes were resuspended in RPMI media (Gibco, #11875093) supplemented with 10% FBS (Gibco), Penicillin/Streptomycin (Gibco), 0.01 μg/ml IL2, 1 μg/ml R848 (Mabtech, #3825-2H) and 50 μM β-mercaptoethanol (Sigma) for ~60 hours stimulation at 37 °C, 5% CO_2_. ELISpot plates (PVDF 96-well plates, Millipore, #MSIPS4510) were coated overnight with 15 μg/ml antigen diluted in PBS, followed by careful washing and blocking using RPMI + 10% FBS. Live splenocytes were counted and the cell number was adjusted to 1×10^7^ cells/ml. Serial dilutions of splenocytes were plated in duplicates and incubated overnight with coated plates. After several wash steps with PBS buffer, plates were incubated for two hours with biotinylated anti-mouse total IgG (Mabtech, # 3825-6-250) in PBS, followed by incubation with streptavidin-conjugated to HRP (Mabtech, #3310-9) for one hour. Spots were revealed using tetramethylbenzidine (TMB, Mabtech, #3651-10) and counted with an automatic reader (Bioreader 2000; BioSys GmbH). Results were represented as number of spots per 10^6^ splenocytes.

### RSV neutralization assay

The RSV A2 strain carrying a luciferase gene (RSV-Luc) was a kind gift of Marie-Anne Rameix-Welti, UFR des Sciences et de la Santé, Paris. Hep2 cells were seeded in Corning 96-well tissue culture plates (Sigma, #CLS3595) at a density of 40,000 cells/well in 100 μl of Minimum Essential Medium (MEM, Gibco, #11095-080) supplemented with 10% FBS (Gibco, 10500-084), L-glutamine 2 mM (Gibco, #25030-081) and penicillin-streptomycin (Gibco, #15140-122), and grown overnight at 37 °C with 5% CO2.

Sera were heat-inactivated for 30 minutes at 56 °C. Serial two-fold dilutions were prepared in an untreated 96-well plate using MEM without phenol red (M0, Life Technologies, #51200-038) containing 2mM L-glutamine, penicillin + streptomycin, and mixed with 800 pfu/well RSV-Luc (corresponding to a final MOI of 0.01). After incubating diluted sera and virus for one hour at 37 °C, growth media was removed from the Hep2 cell layer and 100 μl/well of the serum-virus mixture added. After 48 hours, cells were lysed in 100 μl buffer containing 32 mM Tris pH 7.9, 10 mM MgCl_2_, 1.25% Triton X-100, 18.75% glycerol and 1mM DTT.50 μl lysate were transferred to a 96-well plate with white background (Sigma, # CLS3912). 50 μl of lysis buffer supplemented with 1 μg/ml luciferin (Sigma, #L-6882) and 2 mM ATP (Sigma, #A3377) were added to each well immediately before reading luminescence signal on a Tecan Infinite 500 plate reader.

On each plate, a Palivizumab dilution series was included to ensure comparability of neutralization data. In our assay, we determined IC_50_ values for Palivizumab of 0.32 μg/ml, which is similar to what other groups have reported (*40*). The neutralization curve was plotted and fitted using the GraphPad variable slope fitting model, weighted by 1/Y^2^.

### Sera fractionation

400 μl of streptavidin agarose beads (Thermo Scientific, #20347) were pelleted at 13,000 rpm for 2 minutes in a table top centrifuge and washed with phosphate buffered saline (PBS).200 μg of biotinylated site II peptide were incubated for 2 hours at room temperature to allow coupling of biotinylated peptide to streptavidin beads. Beads were washed three times with 1 ml PBS to remove excess of peptide and resuspended to a total volume of 500 μl bead slurry. Mouse sera from the same cohort (n=10) were pooled (4 μl each, 40 μl total) in a total volume of 200 μl PBS, and 90 μl diluted sera were mixed with 150 μl of bead slurry, followed by an overnight incubation at 4 °C. Beads were pelleted by centrifugation and the supernatant carefully removed by pipetting. Beads were then washed twice with 200 μl PBS and the wash fractions were discarded. To elute site II-specific antibodies, beads were resuspended in 200 μl elution buffer (0.1 M glycine, pH 2.7) and incubated for 1 minute before centrifugation. Supernatant was removed, neutralized with 40 μl neutralization buffer (1 M Tris pH 7.5, 300 mM NaCl), and stored at −20 °C for subsequent testing for RSV neutralization. As a control, unconjugated streptavidin was used for each sample to account for nonspecific binding.

### Next-generation antibody repertoire sequencing (NGS)

#### RNA isolation

Mouse bone marrow was isolated from femurs and re-suspended in 1.5 ml Trizol (Life Technologies, #15596) and stored at −80°C until further processing. RNA extraction was performed using the PureLink RNA Mini Kit (Life Technologies, #12183018A) following the manufacturer guidelines.

#### Antibody sequencing library preparation

Library preparation for antibody variable heavy chain regions was performed using a protocol that incorporates unique molecular identifier (UID) tagging, as previously described in Khan *et al*. (*67*). Briefly, first-strand cDNA synthesis was performed by using Maxima reverse transcriptase (Life Technologies, #EP0742) following the manufacturer instructions, using 5 μg RNA with 20 pmol of IgG gene-specific primers (binding IgG1, IgG2a, IgG2b, IgG2c, and IgG3) with an overhang of a reverse UID (RID). After cDNA synthesis, samples were subjected to a left-hand sided SPRIselect bead (Beckman Coulter, #B23318) cleanup at 0.8X. Quantification of target-specific cDNA by a digital droplet (dd)PCR assay allowed exact input of 135000 copies into the next PCR step. Reaction mixtures contained a forward multiplex primer set that was specific for variable heavy region framework 1 and possessed forward UID (FID), a 3’ Illumina adapter specific reverse primer, and 1X KAPA HIFI HotStart Uracil+ ReadyMix (KAPA Biosystems, #KK2802). PCR reactions were then left-hand side SPRIselect bead cleaned as before and quantified using ddPCR assay. Finally, an Illumina adaptor-extension PCR step was carried out using 820000 copies of the previous PCR product. Following 2nd-step adaptor-extension PCR, reactions were cleaned using a double-sided SPRIselect bead cleanup process (0.5X-0.8X) and eluted in TE buffer.

#### NGS with Illumina MiSeq (2 × 300 bp)

After library preparation, individual NGS libraries were characterized for quality and quantified by capillary electrophoresis using a Fragment Analyzer (Advanced Analytical DNF-473 Standard Sensitivity). Samples were then pooled and NGS was performed on the Illumina MiSeq platform with a MiSeq Reagent Kit V3, 2×300bp paired-end (Illumina, #MS-102-3003), using an input concentration of 10 pM with 10% PhiX.

#### Error and bias correction

Error and bias correction was performed using molecular amplification fingerprinting pipeline, as previously described (*67, 68*).

##### 1) Bioinformatic preprocessing

Paired-end FASTQ files obtained from Illumina MiSeq were imported into CLC Genomics Workbench 10 on the ETH Zurich Euler High Performance Computing (HPC) cluster. A preprocessing workflow was run containing the following steps: trimming of low quality reads, merging of paired-end reads, removal of sequences not aligning to mouse IGH constant sequences, and length filtering.

##### 2) Error correction by consensus building

After pre-processing all datasets were downsampled to contain the same amount of sequencing reads as the dataset with the lowest overall number of reads (361749 sequencing reads). For error correction, a custom Python script was used to perform consensus building on the sequences for which at least three reads per UID were required. VDJ annotation and frequency calculation was then performed by our inhouse aligner (*67, 68*). The complete error-correction and alignment pipeline is available under https://gitlab.ethz.ch/reddy/MAF.

#### Sequence analysis and data visualization

Data analysis was done by customized scripts in R. For the identification of clonotypes hierarchical clustering (*68*) was utilized to group CDR3 sequences together. The following parameters were used: identical IGHV and IGHJ gene segment usage, identical CDR3 length, and at least 80% CDR3 amino acid similarity to one other sequence in the given clonotype (single linkage). The overlap of clonotypes between both cohorts was analyzed by extracting the 20 most expanded clonotypes from each cohort and visualizing their size, occurrence, and Vgene usage by a circos plot using R software circlize (*69*). CDR3 sequence similarities between overlapping clonotypes were represented graphically with the R software motifStack (*70*). All scripts are available upon request.

## Supporting information

## Data availability

Data supporting the findings of this study are available within the article and its Supplementary Information, or are available from the authors upon request.

## Contributions

F.S. and B.E.C. designed experiments. M.G. expressed and purified RSVN and NRM fusion protein. F.S., C.Y., S.S.V and P.C performed ELISAs. F.S. designed and performed SPR competition assay. F.S., C.Y and P.C performed ELISpot experiments, S.S.V, P.C and S.R performed mouse immunizations. L.C and S.F performed next generation sequencing analysis. F.S., J.B.and P.G. designed FFLM. F.S., P.C and M.C. performed RSV neutralization assay. F.S. and B.E.C wrote the paper. All authors commented on the manuscript.

## Acknowledgements

We thank Stefan Kunz for helpful discussions and critical reading of the manuscript. This work was supported by the Swiss initiative for systems biology (SystemsX.ch), the European Research Council (Starting grant - 716058) and the Swiss National Science Foundation.

