## Supplementary material for "Boosting subdominant neutralizing antibody responses with a computationally designed epitope-focused immunogen"

**Fig. S1**

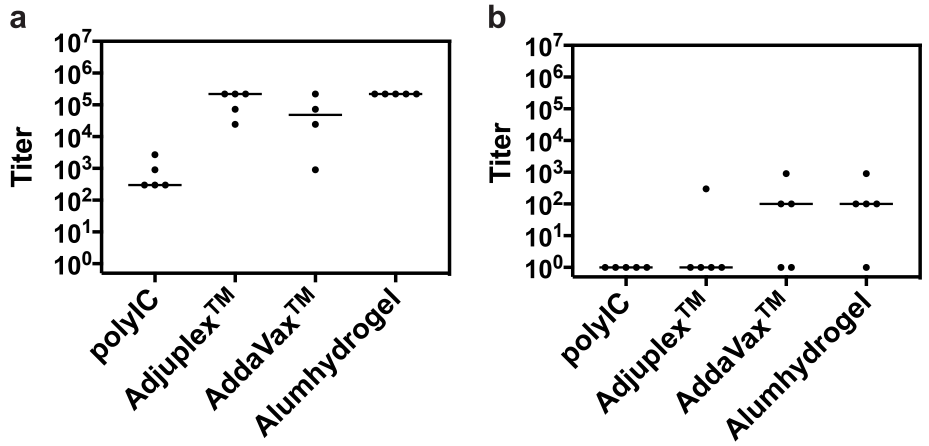


**Supplementary Fig. 1: Adjuvant screen for FFL_001 immunogen. a)** Immunogenicity of FFL_001 formulated in different adjuvants. Serum titers were determined against FFL_001 at day 56 of the immunization protocol. FFL_001 emulsed in Alumhydrogel showed highest overall immunogenicity. **b)** Prefusion RSVF cross-reactivity of FFL_001 immunized mice after three immunizations. 4/5 mice immunized with FFL_001 formulated in Alumhydrogel® showed serum cross-reactivity with prefusion RSVF.

**a**

CLUSTAL O(1.2.4) multiple sequence alignment

```
FFL_001      GSRSDMRKDAERRFDKFVEAAKNKFDKFKAALRKGDIKEERRKDMKKLARKEAEQARRAV     60
FFLM         ASREDMREEADEDFKSFVEAAKDNFNKFKARLRKGKITREHREMMKKLAKQNANKAKEAV     60
             .**.***::*:. *..******::*:**** ****.*..*:*: *****:::*::*:.**

FFL_001      RNRLSELLSKINDMPITNDQKKLMSNDVLKFAAEAEKKIEALAADAEDKFTQ 112
FFLM         RKRLSELLSKINDMPITNDQKKLMSNQVLQFADDAEAEIDQLAADATKEFTG 112
             *:************************:**:** :** :*: ***** .:**
```

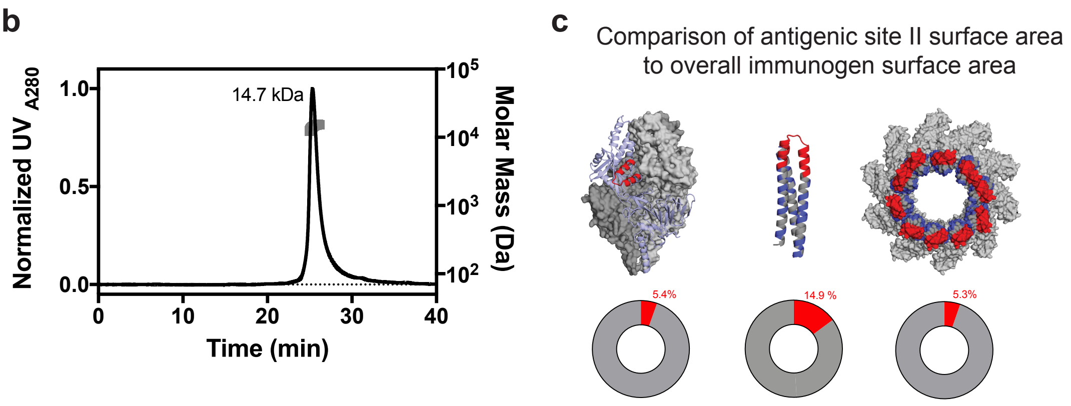


**Supplementary Fig. 2**: **Homology guided resurfacing of FFL_001. a)** Sequence alignment of FFL_001 and FFLM. **b)** Resurfaced variant FFLM is monomeric in solution as assessed by size-exclusion coupled to an on-line multi-angle-light scattering detector. Determined mass in solution is 14.7 kDa ± 3.5 %, which is close to the theoretical molecular weight of 14.4 kDa. **c** Relative surface area of RSVF antigenic site II in prefusion RSVF (PDBID 4JHW), FFL_001 and FFLM models based on crystal structure of RSV N (PDBID: 2WJ8). The Motavizumab epitope is highlighted in red, blue patches indicate sequence changes of FFLM compared to FFL_001, and piecharts show the fraction of antigenic site II surface area compared to overall immunogen surface area. Solvent accessible surface area (SASA) was computed in PyMol in presence and absence of Motavizumab bound to every site II repetition. % SASA of antigenic site II is nearly identical between RSVF and NRM, whereas FFLM monomer shows approximately 3-fold higher surface area of antigenic site II, due to its small size.

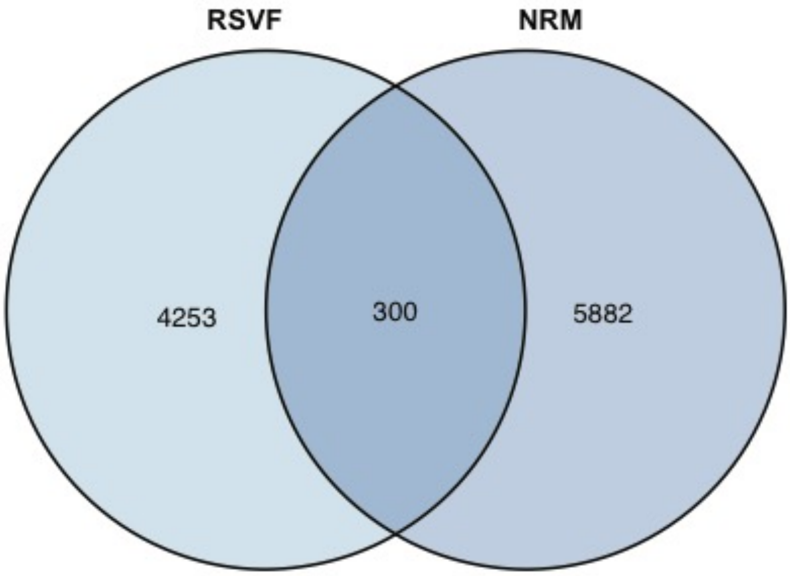


**Supplementary Fig. 3: Overlapping clonotypes obtained from next-generation antibody repertoire sequencing of mice immunized with RSVF or NRM.** When comparing clonotypes, defined as the same VH gene and 80% sequence similarity in the HCDR3, NRM and RSVF immunizations yield 300 overlapping clonotypes.

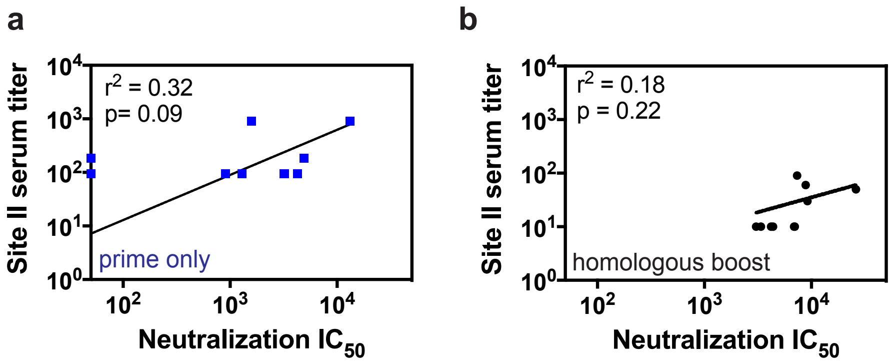


**Supplementary Fig. 4: Correlation of site II specific serum titer (measured by peptide ELISA) with RSV neutralization $IC_{50}$.** Correlation for mouse cohort receiving only a prefusion RSVF priming immunization (a) and the homologous boost cohort (b). Data represent the mean of two independent experiments, each measured in duplicates. Pearson correlation coefficients ($r^2$) and p-values were calculated in GraphPad Prism.

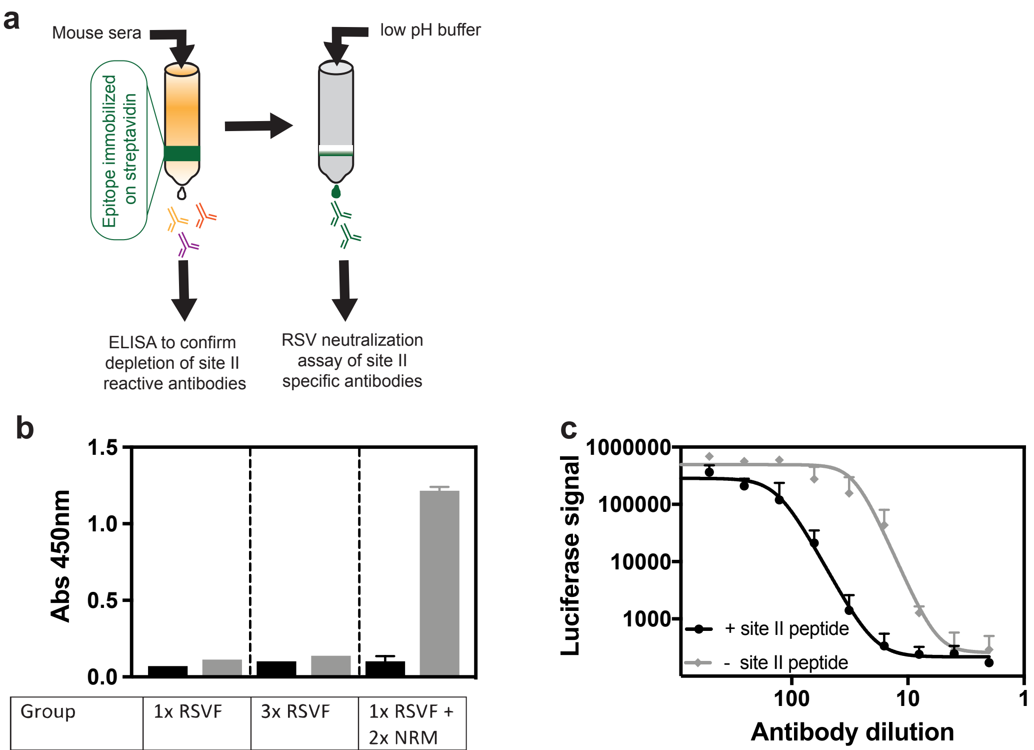


| Group | 1x RSVF | | 3x RSVF | | 1x RSVF + 2x NRM | |
|---|---|---|---|---|---|---|
| Site II peptide conjugated | + | - | + | - | + | - |

**Supplementary Fig. 5: Serum fractionation and enrichment of site II specific antibodies. a)** Experimental scheme. Streptavidin agarose beads were conjugated to biotinylated antigenic site II peptide. As control, unconjugated streptavidin beads were prepared. Sera from ten mice within each cohort was pooled and mixed with conjugated and unconjugated beads. Column flow-through was analyzed by ELISA (b), and eluted site II specific antibodies were analyzed in an RSV neutralization assay (c). **b)** ELISA against antigenic site II peptide of column flow-through. Immunization groups as described in Figure 4 (prime only, homologous boost and heterologous boost). ELISA signal (OD at 450 nm) is shown for column flow-through from serum fractionation shown in (a). Streptavidin beads that were not coupled to antigenic site II were used as controls, and did not deplete site II reactivity in the flow through. Data and error bars presented are averaged from two different experiments. **c)** Example RSV neutralization assay curves from elution fraction, using site II conjugated (black) or unconjugated streptavidin beads (grey). Luciferase signal is plotted on the y-axis, and is a measure for RSV replication as previously reported (*1*). The elution fraction is diluted as indicated on the x-axis. Data shown are from one experiment performed in duplicates.

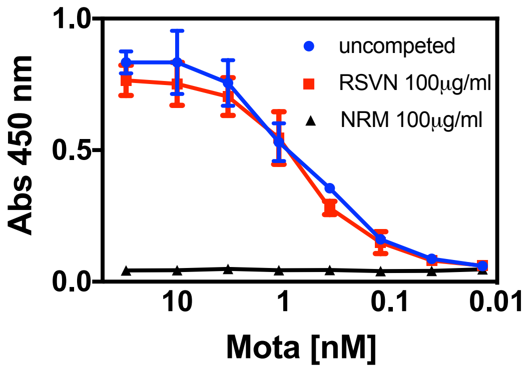


**Supplementary Fig. 6: Competition ELISA with Motavizumab antibody control**. Plates were coated with prefusion RSVF as described in methods. A 3-fold serial dilution of Motavizumab (initial concentration = 30 nanomolar) was prepared in presence of different competitors, either the unconjugated RSV nucleoprotein nanoparticle (RSVN), the epitope scaffold nanoparticle (NRM), or none. Following overnight competition, binding of Motavizumab to RSVF was measured. RSVN competition did not affect RSVF binding of Motavizumab as expected. In contrast, NRM efficiently competed with RSVF for Motavizumab binding at the indicated competitor concentration. Data shown are from one experiment, with error bars derived from technical duplicates.

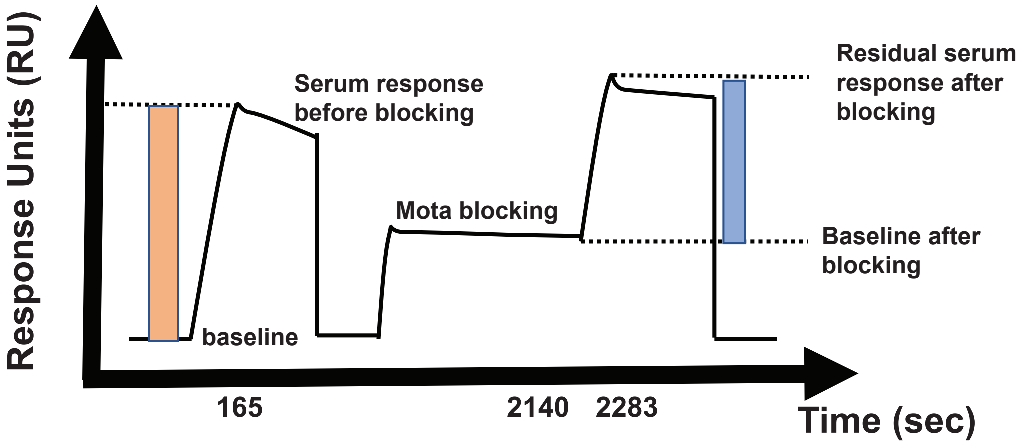


**Supplementary Fig. 7: Schematic representation of the Surface Plasmon Resonance competition assay.** Mouse sera is injected on an antigen coated sensor chip surface to measure initial response (orange). Following regeneration, Motavizumab binding sites are blocked by excess of Motavizumab. Residual serum response is determined on a blocked surface (blue). For data analysis, response units at indicated timepoints are extracted, and % competition is calculated as described in methods and shown in Supplementary Table 1.

**Prime only**

| Timepoint (sec) | mouseID C1 | C2 | C3 | C4 | C5 | S1 | S2 | S3 | S4 | S5 |
|---|---|---|---|---|---|---|---|---|---|---|
| 165 | 33.6 | 39.1 | 36.1 | 24.0 | 10.8 | 45.5 | 23.4 | 11.5 | 18.8 | 36.5 |
| 2140 | 72.2 | 74.9 | 100.4 | 102.2 | 110.2 | 204.5 | 151.6 | 169.3 | 110.8 | 116.4 |
| 2283 | 99.4 | 107.9 | 132.4 | 121.5 | 119.8 | 236.8 | 164.1 | 176.7 | 126.5 | 149.8 |
| **% blocking** | **19.3** | **15.3** | **11.7** | **19.9** | **11.7** | **29.0** | **46.3** | **36.2** | **17.0** | **8.5** |

**Heterologous boost**

| Timepoint (sec) | mouseID A1 | A2 | A3 | A4 | A5 | U1 | U2 | U3 | U4 | U5 |
|---|---|---|---|---|---|---|---|---|---|---|
| 165 | 41.1 | 20.0 | 44.9 | 34.1 | 19.5 | 28.5 | 21.3 | 44.0 | 49.1 | 27.0 |
| 2140 | 131.1 | 92.7 | 95.7 | 101.9 | 97.5 | 281.4 | 191.9 | 203.3 | 215.4 | 157.4 |
| 2283 | 165.5 | 108.1 | 111.5 | 123.3 | 112.1 | 300.2 | 201.9 | 227.0 | 244.4 | 175.2 |
| **% blocking** | **16.5** | **23.2** | **64.8** | **37.1** | **25.5** | **34.1** | **53.1** | **46.0** | **40.8** | **34.1** |

**Homologous boost**

| Timepoint (sec) | mouseID W1 | W2 | W3 | W4 | W5 | RSVF1 | RSVF2 | RSVF3 | RSVF4 | RSVF5 |
|---|---|---|---|---|---|---|---|---|---|---|
| 165 | 54.8 | 75.6 | 42.1 | 52.5 | 51.3 | 65.7 | 62.8 | 75.8 | 54.7 | 66.9 |
| 2140 | 123.8 | 119.2 | 91.4 | 106.0 | 99.4 | 104.8 | 110.4 | 104.2 | 81.5 | 92.0 |
| 2283 | 169.2 | 182.8 | 121.9 | 148.5 | 142.2 | 159.2 | 158.7 | 163.6 | 126.5 | 146.6 |
| **% blocking** | **17.1** | **15.9** | **27.8** | **19.1** | **16.6** | **17.1** | **23.1** | **21.7** | **17.7** | **18.5** |

**Supplementary Table 1: Response units from SPR competition assay for prime boost immunization regimen.** Timepoints as shown in Supplementary Fig. 7.

1. M. A. Rameix-Welti *et al.*, Visualizing the replication of respiratory syncytial virus in cells and in living mice. *Nat Commun* **5**, 5104 (2014).
